# AGOUTI: improving genome assembly and annotation using transcriptome data

**DOI:** 10.1101/033019

**Authors:** Simo V. Zhang, Luting Zhuo, Matthew W. Hahn

## Abstract

**Summary:** Current genome assemblies consist of thousands of contigs. These incomplete and fragmented assemblies lead to errors in gene identification, such that single genes spread across multiple contigs are annotated as separate gene models. We present AGOUTI (Annotated Genome Optimization Using Transcriptome Information), a tool that uses RNA-seq data to simultaneously combine contigs into scaffolds and fragmented gene models into single models. We show that AGOUTI improves both the contiguity of genome assemblies and the accuracy of gene annotation, providing updated versions of each as output.

**Availability:** The software is implemented in python and is available from github.com/svm-zhang/AGOUTI.

**Supplementary information:** Supplementary data are available at *Bioinformatics* online.

## 1 Introduction

Genomes sequenced using short-read, next-generation sequencing technologies are error-filled and fragmented into thousands of small sequences (Alkan et al., 2011; Ye et al., 2011). In addition to a general lack of data about sequence contiguity, one consequence of fragmented genome assemblies is that single genes are placed on multiple contigs or scaffolds, increasing the number of predicted genes (Denton et al., 2014). Such biases can confound inferences about the number of genes within species, and gene gain and loss between species (Han et al., 2013).

Data from expressed genes (i.e. transcriptome or RNA-seq data) has previously been used to combine contigs into scaffolds (e.g. Xue et al., 2013; Mortazavi et al., 2010), acting in effect as a mate-pair library with insert size equivalent to intron length. Although this use results in an improvement in contiguity, it does not generally decrease the number of incorrectly predicted genes. This is because contigs within scaffolds are connected by gaps, and gene prediction programs cannot predict across gaps of even moderate length. However, we previously showed that RNA-seq can also be used to reduce the number of gene models split apart by fragmented assembles because it contains information about connections between exons in a single gene (Denton et al., 2014).

Here we combine these two uses of transcriptome data into a single lightweight program that we call AGOUTI. As with other scaffolders based on RNA-seq (Xue et al., 2013; Mortazavi et al., 2010), AGOUTI brings together contigs into scaffolds, yielding a more contiguous assembly. It does this with an algorithm similar to the one used in RNAPATH (Mortazavi et al., 2010), but with additional denoising steps and constraints that ensure greater accuracy. AGOUTI also simultaneously updates gene annotations by connecting predictions from multiple contigs, significantly reducing the number of gene models initially predicted from draft assemblies. We are not aware of other annotation software that has these features.

## 2 Methods

An overview of AGOUTI is given in **Supplementary Figure 1**. The method takes three inputs: an initial genome assembly in FASTA format, paired-end RNA-seq reads mapped against this assembly in BAM format, and gene predictions from the initial assembly in GFF3 format. The output of AGOUTI will be an updated genome assembly file (in FASTA format) and an updated set of gene predictions (in GFF3 format).

AGOUTI carries out scaffolding by first constructing an edge-weighted adjacency graph made up of contigs (vertices) and the supporting joining-pairs between them (edges). AGOUTI then denoises the graph by removing erroneous joining-pairs based on the presence of intervening genes and read orientation (see **Supplementary Materials; Supplementary Figures 2 and 3**). Our results show that traversing a graph without noisy edges places many more contigs together accurately, generating an assembly with higher contiguity (see Results). AGOUTI traverses the graph from each vertex and follows the highest-weighted edges until no further extension can be made (**Supplementary Materials; Supplementary Figure 4A**). Each walk gives a scaffolding path, where the shortest such path includes only two contigs. AGOUTI then reconciles each path using constraints imposed by the constituent gene models (**Supplementary Materials; Supplementary Figure 5**). For each reconciled path, AGOUTI joins contigs into scaffolds, separating them by a gap of length defined by the user (1-kb by default). Contigs are reverse-complemented whenever needed.

AGOUTI also updates gene models according to the new assembly. For each pair of contigs within a scaffold, AGOUTI merges the two gene models from which the connection was made. The gene-merge assigns a new gene ID with prefix “AGOUTI” in the output, combines exons, and converts coordinates to the new scaffold system. If contigs are reverse-complemented, all gene models on that contig will be reversed accordingly in the output annotation.

## 3 Results

### 3.1 Data

To evaluate the performance of AGOUTI, we randomly fragmented the genome of the N2 strain of *Caenorhabditis elegans* (version WS246; The *C. elegans* Sequencing Consortium 1998; Stein et al., 2001) into six assemblies with varying numbers of contigs (**Supplementary Table 1**). For each fragmented assembly, we performed gene prediction using AUGUSTUS (Stanke et al., 2003). We found that assemblies with larger numbers of contigs had increased numbers of predicted gene models (square boxes in **Fig. 1**), consistent with results reported by Denton et al. 2014. We used a single RNA-seq dataset from the same strain of *C. elegans* at the early embryo stage (SRR316753, SRR317082, and SRR350977). We mapped these reads against each of our fragmented assemblies using BWA-mem (Li 2013), and used the mapping results (Li et al., 2009) along with the predicted gene models, as inputs to AGOUTI.

**Fig. 1.**
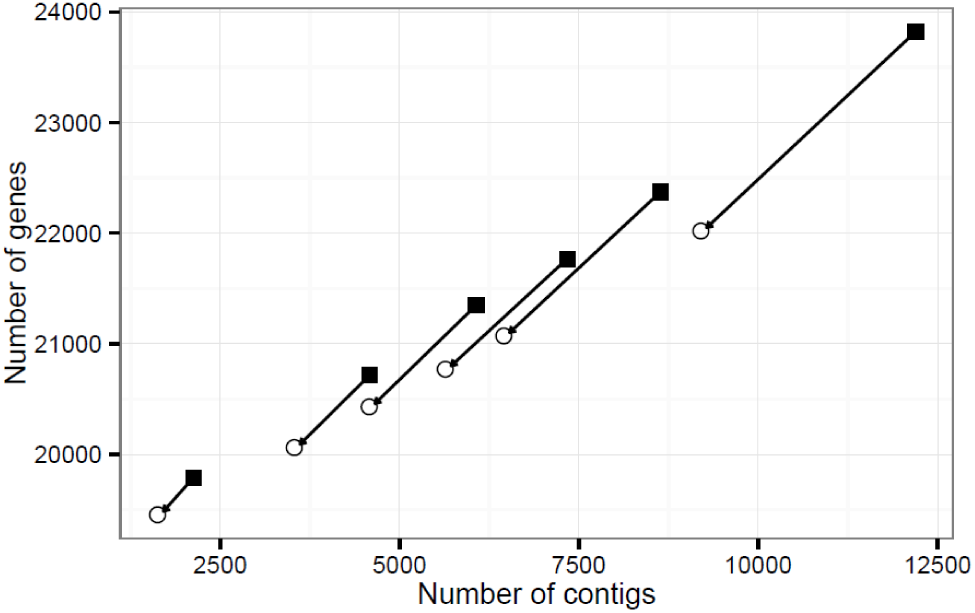
Performance of AGOUTI scaffolding. AGOUTI was able to scaffold hundreds to thousands of contigs, and significantly reduced the number of gene models. The least contiguous assembly has the largest number of contigs scaffolded and the largest number of gene models reduced. The x-axis represents each simulated assembly in terms of number of contigs. The leftmost gives the most contiguous assembly, while the one on the far right is the least contiguous assembly. The number of gene models is on the y-axis. The square boxes and open circles give the number of gene models before and after the scaffolding, respectively.

### 3.2 Evaluation

We evaluated the performance of AGOUTI on the six assemblies using default settings. AGOUTI was able to scaffold hundreds to thousands of contigs (open circles in **Fig. 1**), yielding higher scaffold N50 values (**Supplementary Table 1**). The most fragmented assembly had the largest number of contigs joined into scaffolds and the largest number of gene models reduced (**Fig. 1**). We checked the accuracy of contigs placed within each scaffold by comparing the output of AGOUTI to the N2 reference assembly. Across our simulated assemblies, AGOUTI achieved high accuracy by putting at least 99.95% of contig pairs in the correct order (using a minimum of 5 joining-pairs, **Supplementary Table 3**). We found zero pairs of contigs across our six assemblies that were incorrectly ordered (intra-chromosomal error; **Supplementary Table 3**), but did find two cases where two contigs coming from different chromosomes were placed together (inter-chromosomal error; **Supplementary Table 3**). We compared results using AGOUTI to results obtained from RNAPATH, across a range of different input values (Supplementary Materials). Across all conditions, AGOUTI found more connections than RNAPATH (compare **Supplementary Table 1** to **Supplementary Table 2**) and had fewer errors (compare **Supplementary Table 3** to **Supplementary Table 4**).

We also investigated whether the connections between contigs made by AGOUTI accurately reflect the existence of underlying genes. Specifically, we asked for each contig pair whether the joining-pairs used for scaffolding were mapped to two exons of a single gene for each contig pair (**Supplementary Figure 6**). We used the gene annotation of the same version as the reference N2 genome to evaluate these connections (Stein et al. 2001). Within each assembly, approximately 95% of genes joined by AGOUTI connected two exons of the same annotated gene (using a minimum of 5 joining-pairs; **Supplementary Figure 6A**, **Supplementary Table 5**). Interestingly, among the rest of the contig pairs, some connected an exon on one contig with an unannotated exon on the other contig (Case 1, **Supplementary Figure 6B**). Another class of genes merged by joining-pairs had mappings to two different genes (Case 2, **Supplementary Figure 6C**). This may indicate that these two genes should be merged into one. In the last scenario, both ends of the joining-pairs failed to map to any known genes on both contigs, suggesting a potential novel gene (Case 3, **Supplementary Figure 6D**). For consecutive pairs of contigs (i.e. pairs that are physically next to each other on a chromosome), we considered these interesting cases to be a bonus feature of AGOUTI, and did not count them as false positives; the number of each type is listed in **Supplementary Table 5**.

## 4 Conclusion

AGOUTI is a powerful and effective scaffolder, and unlike most scaffolders is expected to become more effective in larger genomes because of the commensurate increase in intron length. AGOUTI is able to scaffold thousands of contigs while simultaneously reducing the number of gene models by several thousand. It therefore makes it easy to improve both genome assemblies and genome annotations.

## Acknowledgements

This work was supported by National Science Foundation grant DEB-1249633 to M.W.H.

*Conflict of Interest:* none declared.

